# Benzo[a]pyrene degrading *Bacillus altitudinis* AR19 isolated from Digboi oil refinery soil (India)

**DOI:** 10.1101/2025.09.16.676557

**Authors:** Abhisek Dasgupta, Bardwi Narzary, Riya Singh

## Abstract

Benzo[a]pyrene (B[a]P) is a five-ring polycyclic aromatic hydrocarbon (PAH) recognized as a potent carcinogen and persistent environmental pollutant, highly resistant to microbial degradation. In this study, a B[a]P-degrading and biosurfactant producing bacterium, *Bacillus altitudinis AR19*, was isolated from the Digboi oil refinery site in Assam, India. The isolate was screened using the spray plate technique, and its morphological features and cell viability were examined by scanning electron microscopy (SEM) and flow cytometry. Biosurfactant production was confirmed through drop collapse, oil spreading, and emulsification assays. Taxonomic identity was established using average nucleotide identity (ANI), and comparative pangenome analysis was performed among *B. altitudinis* (Taxonomy ID: 1178544), *Bacillus pumilus* (Taxonomy ID: 1408), *Mycolicibacterium vanbaalenii* (Taxonomy ID: 110539), and *B. altitudinis* AR19. The results highlight that several mechanisms including osmolyte accumulation, biosurfactant production, and mechanosensitive ion channels play key roles in enabling B. altitudinis AR19 to tolerate and degrade B[a]P. This study demonstrates the potential of B. altitudinis AR19 as a promising candidate for bioremediation of PAH-contaminated environments.

**Importance of the work:** This manuscript describes the morphological and genomic characterization of *Bacillus altitudinis* AR19, a bacterium capable of degrading benzo[a]pyrene. Benzo[a]pyrene is a five-ring polycyclic aromatic hydrocarbon that is highly recalcitrant to bacterial remediation, making this work important for advancing bioremediation strategies.

## Introduction

*Bacillus altitudinis* is a ubiquitous bacterium with diverse properties, ranging from soil bioremediation to plant growth promotion (1,2). This species was first identified in UV-stressed air samples collected from the stratosphere (3). It is known to tolerate high salt concentrations (4), degrade xylan-based polysaccharides (5), and even exhibit herbicidal activity (6).

Polycyclic aromatic hydrocarbons (PAHs) are persistent environmental pollutants that pose serious risks to both ecosystems and human health (7). Due to their lipophilic nature, PAHs are readily absorbed in the gastrointestinal tract of animals (8). Among them, benzo[a]pyrene (B[a]P), a high–molecular-weight PAH composed of five fused benzene rings, is of particular concern. It is primarily released into the environment through the incomplete combustion of fossil fuels and the pyrolysis of organic matter(9). B[a]P is highly toxic, and its metabolic activation leads to the formation of DNA adducts that can induce mutations and carcinogenesis (9). However, its hydrophobicity makes B[a]P highly recalcitrant to microbial degradation (10), and only a limited number of bacterial species capable of degrading it have been reported.

This study reports the isolation of a bacterial strain, bacillus altitudinis AR19, from oil-contaminated soil of the Digboi oil refinery in Assam, India. The strain was screened for its ability to degrade benzo[a]pyrene (B[a]P), and the effects of B[a]P exposure were analyzed using Scanning Electron Microscopy (SEM) and Flow Cytometry. Biosurfactant production by AR19 was evaluated against crude oil. Whole-genome sequencing confirmed the species identity, and a pangenome analysis was performed in comparison with bacillus altitudinis (Taxonomy ID: 1178544), bacillus pumilus (Taxonomy ID: 1408), and mycolicibacterium vanbaalenii (Taxonomy ID: 110539). The overall objective of this study was to isolate and characterize a B[a]P degrading bacterium from oil contaminated soil.

## Materials and Methods

### Isolation and screening of B[a]P degrading bacteria

Soil samples were collected from the Digboi oil refinery (Assam, India). Bacterial strains were isolated using the serial dilution technique (11). Screening for benzo[a]pyrene (B[a]P)-degrading bacteria was performed by the modified spray plate method(12). A stock solution of B[a]P (100 µg/ml) was prepared in dichloromethane, and 10 µl of this solution was evenly sprayed onto the agar plates to assess bacterial growth and degradation activity.

### Flow Cytometry and SEM analysis of the positive isolate

The positive isolate was cultured in liquid medium containing B[a]P, and a time-dependent study was conducted to assess cell viability using flow cytometry. Cells were stained with the LIVE/DEAD™ BacLight™ Bacterial Viability and Counting Kit (Thermo Fisher Scientific, Cat. No. L34856). Scanning Electron Microscopy (SEM) was employed to compare the morphology of B[a]P-treated AR19 cells with untreated controls.

### Screening for biosurfactant production

Bacillus altitudinis AR19 was cultured in nutrient broth, and cells were harvested at the logarithmic growth phase. The culture was centrifuged at 8,000 rpm for 10 minutes, and the resulting cell-free supernatant was subjected to drop-collapse, oil-spreading, and emulsification capacity assays (13).

### Genomic DNA extraction and sequencing

Genomic DNA extraction and sequencing were performed for strain AR19, as described previously in our published manuscript (14)

### Species detection and pangenome analysis

For species identification and pangenome analysis, phylogenetic classification of AR19 was carried out using the GTDB-Tk tool (15). Pangenome analysis was conducted with OrthoMCL(16), comparing the query strain with bacillus altitudinis (Taxonomy ID: 1178544), bacillus pumilus (Taxonomy ID: 1408), and mycolicibacterium vanbaalenii (Taxonomy ID: 110539).

## Results and Discussion

From the soil samples collected at the Digboi oil refinery, a total of 10 bacterial strains were isolated, of which only one strain demonstrated the ability to utilize benzo[a]pyrene (B[a]P). This strain was designated AR19. Screening was performed using a modified spray plate technique (12), in which the B[a]P solution was sprayed directly onto the agar surface without the use of a filter membrane.

Flow cytometry analysis (Figure 1) revealed that after 7 days of incubation in B[a]P-containing medium, only 57.54% of AR19 cells remained viable. Cell viability was assessed in a closed system, where microbial biomass accumulation altered the nutrient balance of the medium. Specifically, once the proportion of dead cells exceeded 50%, the C:N ratio shifted due to an increase in carbon content from lysed biomass and a relative decrease in available nitrogen, which further reduced cell viability.

**Figure 1.**
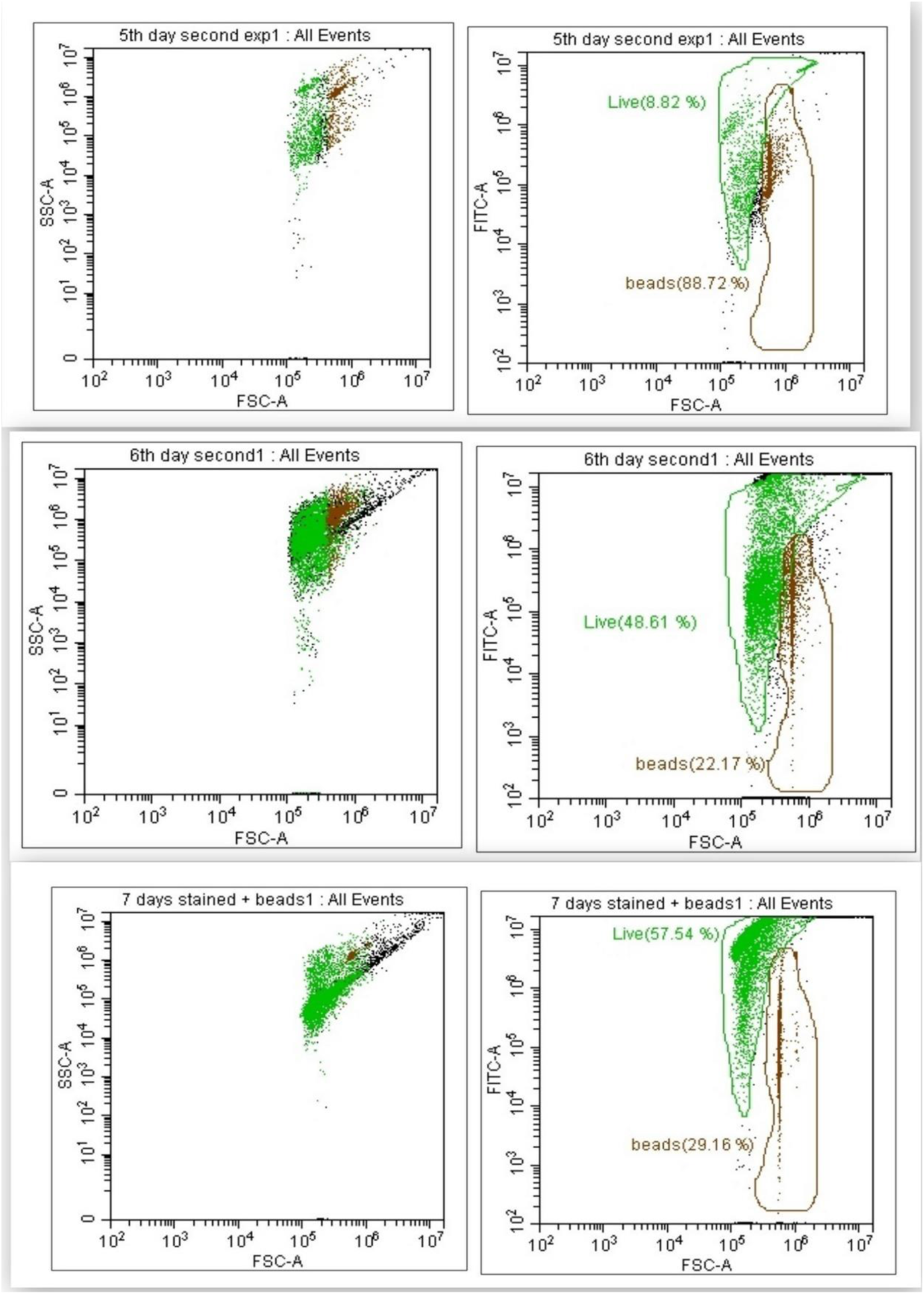
Flow cytometry dot plot depicting the viability of *Bacillus altitudinis AR19 cells*, with the highest percentage (57.54%) observed after 7 days of incubation.

Scanning electron microscopy (SEM) images (Figure 2) of AR19 cells after 24 h and 72 h of incubation showed marked differences in cell morphology. With prolonged exposure to B[a]P, the cells appeared swollen, suggesting that AR19 experienced osmotic stress under these conditions.Biosurfactant activity of strain AR19 was confirmed through multiple assays. In the drop collapse test (Figure 3), all three replicates showed an increase in drop diameter from the initial 8 mm to 11–13 mm after 1 h, indicating a reduction in surface tension. In contrast, the control (distilled water) maintained a constant diameter of 6 mm with no collapse. In the oil displacement assay, clear zones of 15–36 mm were observed across replicates, further confirming biosurfactant production (Figure3). The emulsification index (E24) ranged from 6.66% to 22.22%, with the highest activity recorded in replicate 3 (Figure 3). Together, these results demonstrate the ability of AR19 to produce effective biosurfactants with surface-active properties.

**Figure 2.**
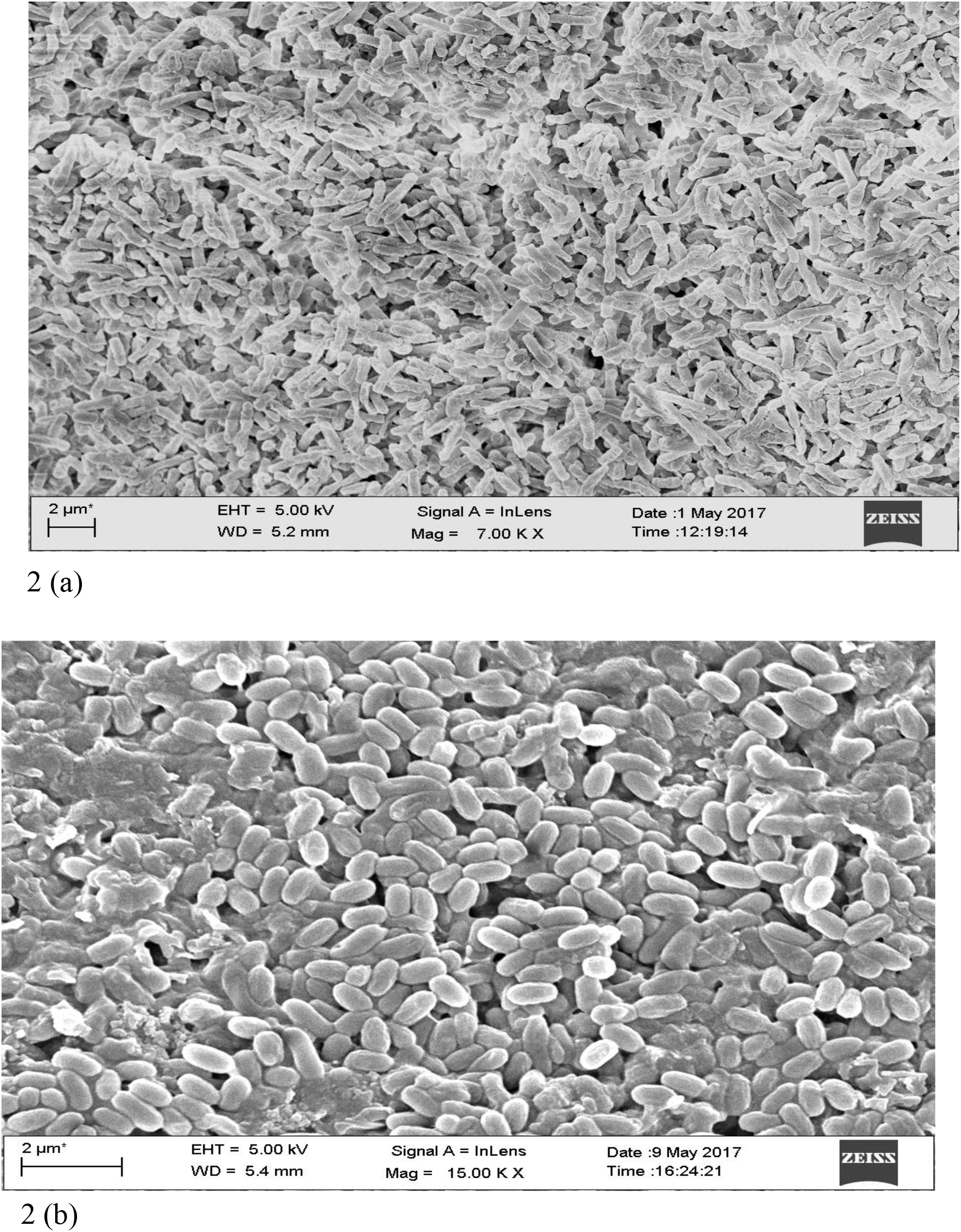
Scanning electron micrographs of *Bacillus altitudinis AR19* at (a) 24 h and (b) 72 h of incubation with benzo[a]pyrene. Cells exhibited swelling and a rounded morphology after 72 h.

**Figure 3.**
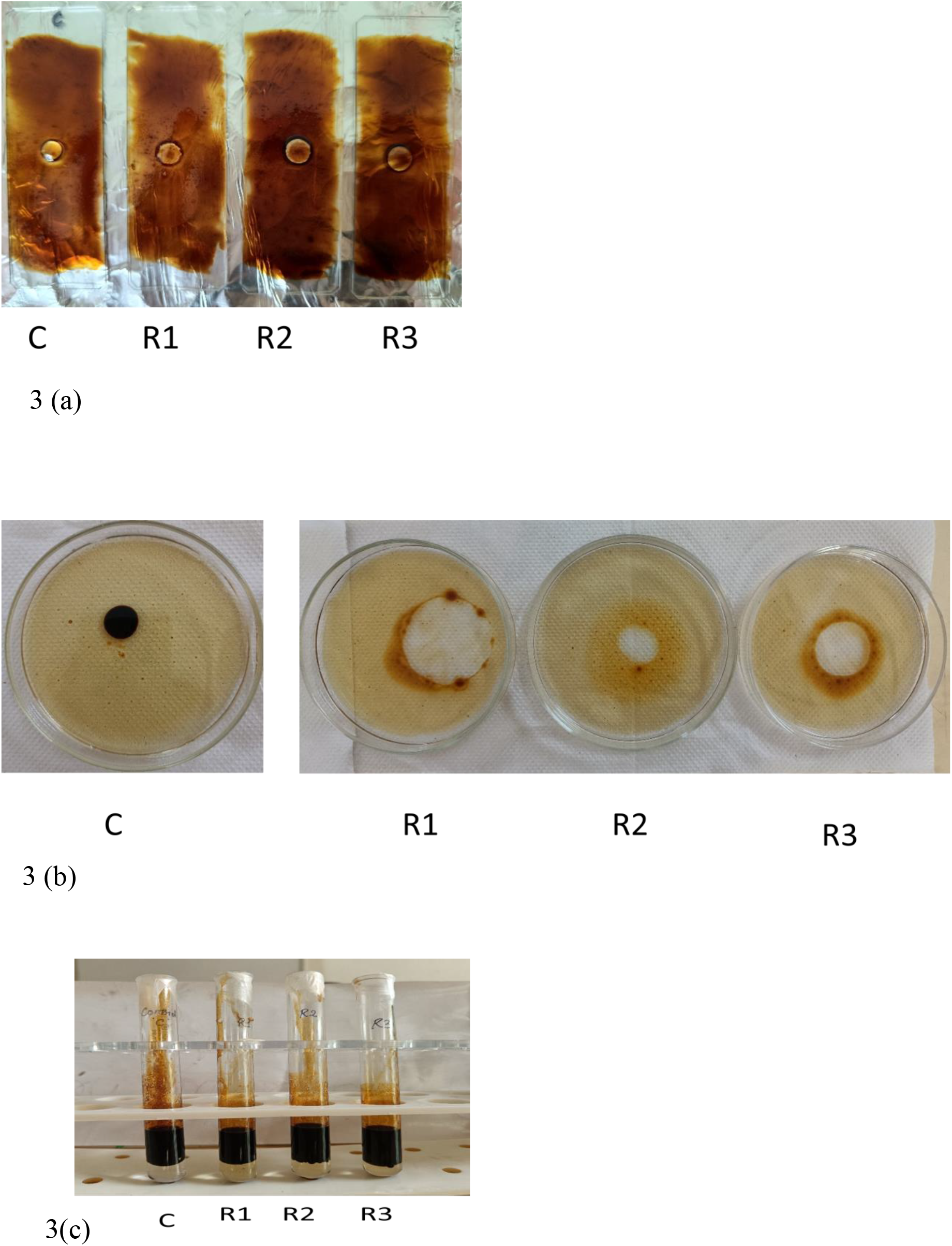
Biosurfactant screening of Bacillus altitudinis AR19, in the (a) drop collapse test it was oserved all the replicates (R1-R3) indicated a reduction in surface tension after 1 hour which was not the case of control (C) (b) Oil displacement assay indicating clear zone in the replicates (R1-R3) as compared to control (C) (c) The emulsification index assay showing moderate to good emulsifying activity in the samples.

Phylogenetic analysis (Figure 4) showed that the AR19 genome shared a branch with *Bacillus pumilus*, indicating a close relationship and low genetic divergence. Additionally, its taxonomic classification by the ANI method revealed its closest reference as *Bacillus altitudinis*. Several protein-encoding genes that provide resistance to hydrocarbon stress were annotated in the genome. Specifically, genes involved in iron uptake, sensing oxygen gradients, biofilm formation, complex carbohydrate utilization, UV protection, PAH metabolism, and functioning as antiporters and symporters were considered important for its survival in a hydrocarbon-enriched environment (Table 1). Hydrophobic substances cause cell toxicity by inducing water stress (17). The positive isolate possessed large conductance mechanosensitive ion channels (MscL) and an OpuA operon in its genome, so it might be possible that the initial response to osmotic stress is mediated by these genes. The MscL channel opens when the lipid bilayer stretches due to water tension, helping the bacterium regulate changes in intracellular osmotic pressure (18). This is supported by the observed swelling of the positive isolate’s cells during incubation with B[a]P, as seen in the SEM micrograph (Figure 2). The OpuA operon is expressed under osmotic stress. In Bacillus species, proline is the principal osmolyte that protects the cell against a hypersaline environment (19). Because the synthesis of proline takes a very long time to reach an effective defensive level (20), the Bacillus sp. synthesizes glycine and betaine from choline to protect its cell during the initial stage of osmotic stress. These compounds are taken up directly from the environment with the help of the OpuA transporter (21).

**Table 1.**
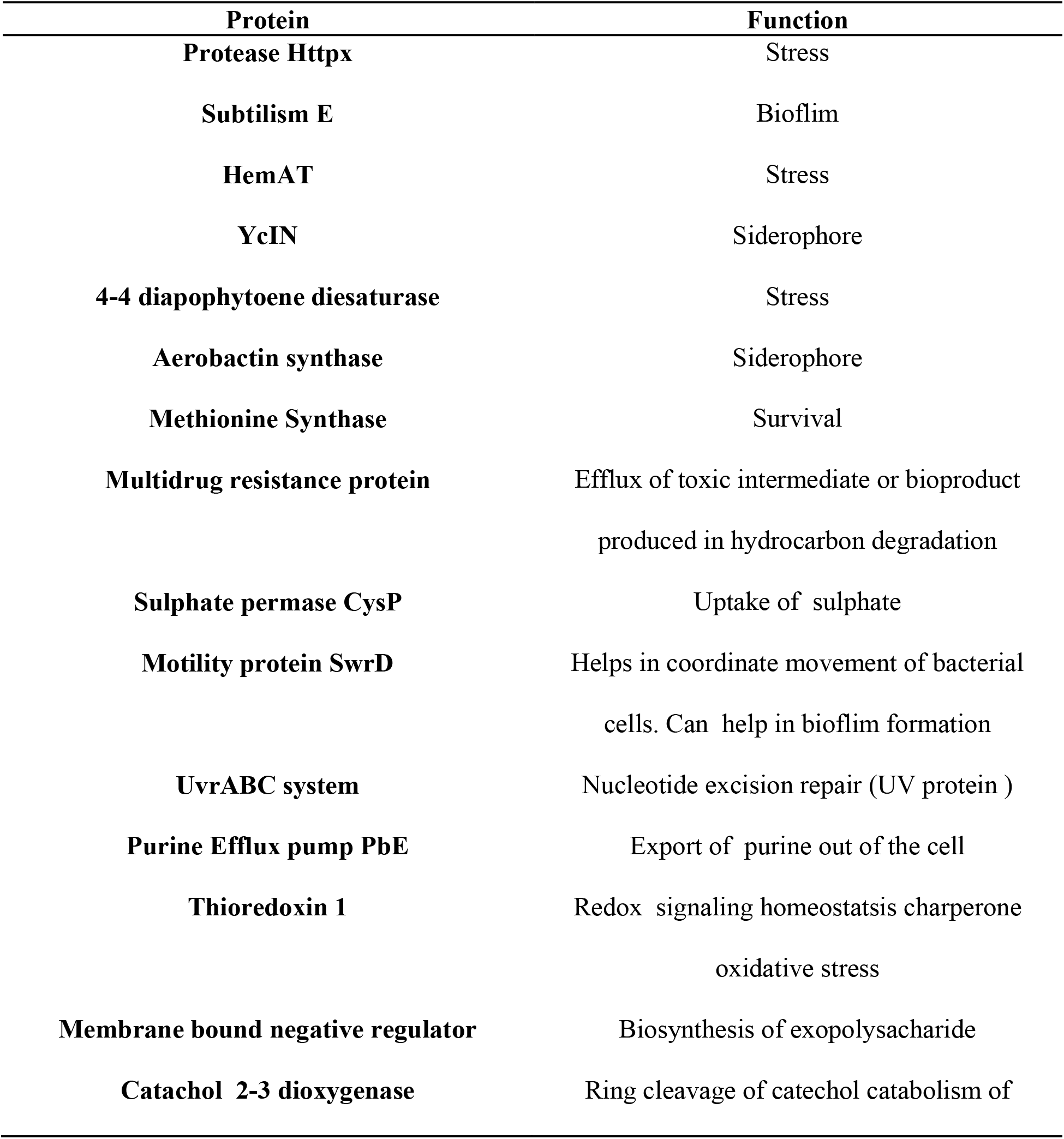
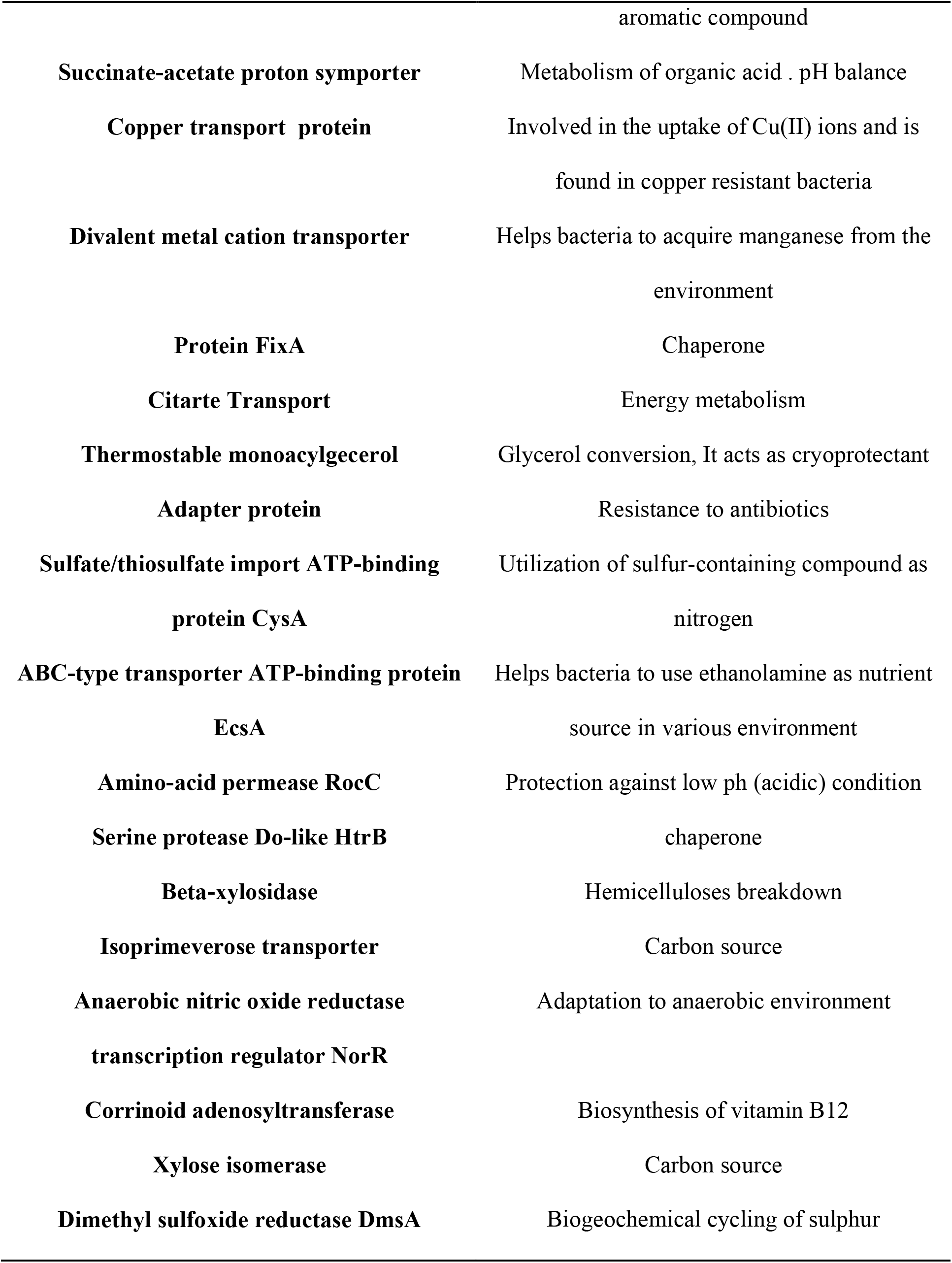
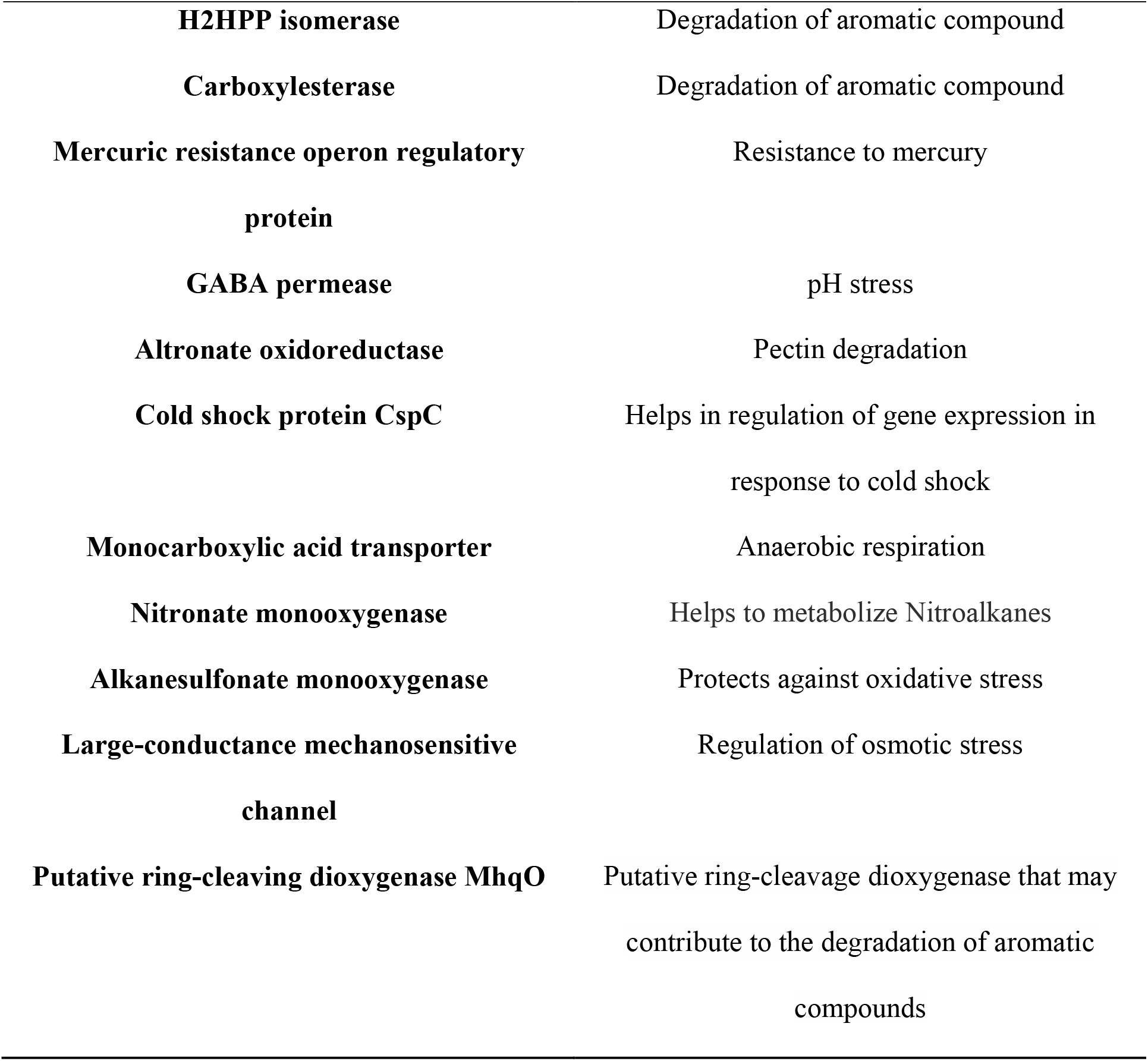
Annotated proteins of *bacillus altitudinis* AR19 and their proposed function.

**Figure 4.**
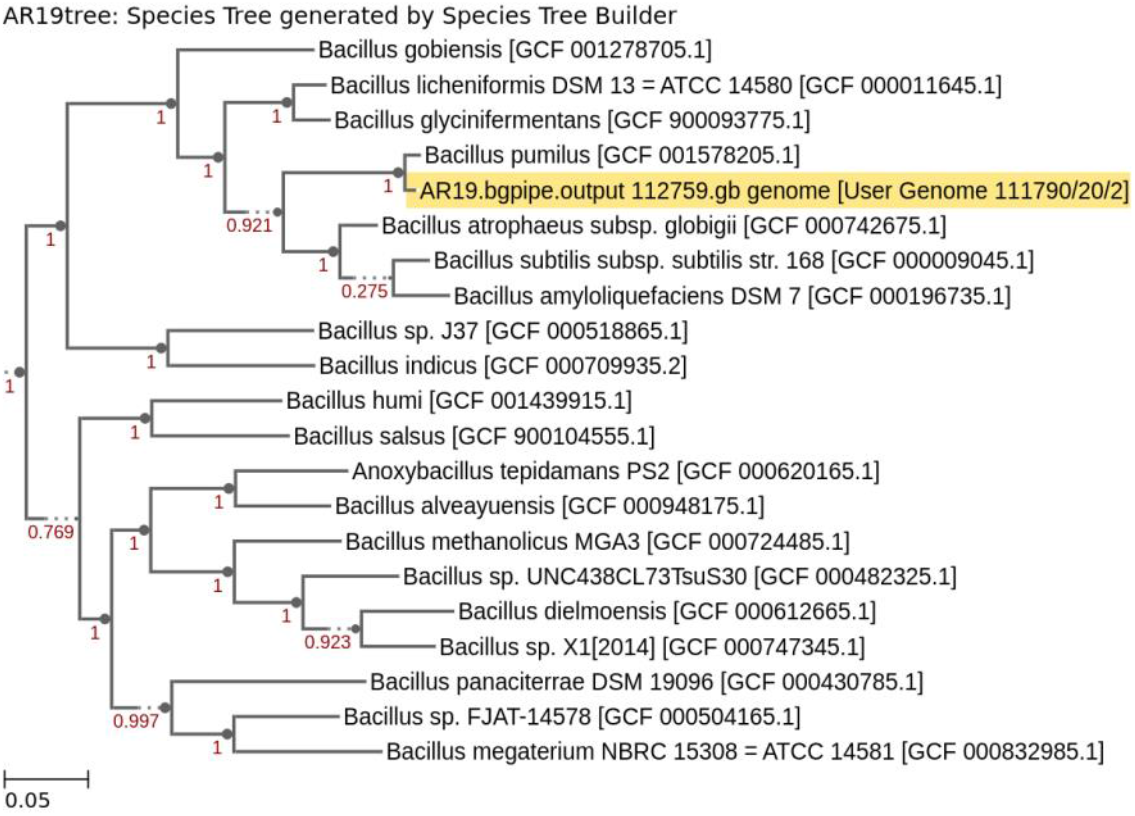
Species tree showing the phylogenetic placement of Bacillus altitudinis AR19. The strain clustered with *bacillus pumilus*, indicating a close evolutionary relationship.

A possible degradation pathway for B[a]P was proposed from the genome annotation. Surfactin and biofilm play a crucial role in making B[a]P accessible to the positive isolate. Surfactin, a biosurfactant, has a good emulsifying property (22) and the biofilm helps to stabilize the droplets of B[a]P which have been dispersed in the aqueous phase (23) Through these two mechanisms, the isolate adsorbs B[a]P from the aqueous media. The isolate uses an extradiol ring cleavage mechanism to degrade B[a]P into a carbon source. The putative ring-cleaving dioxygenase, MhqA, starts the cleaving process using molecular O_2_.The derivatives are further processed by Catechol-2,3-dioxygenase, encoded by the catE gene, into 2-hydroxymuconate semialdehyde (24). It then undergoes further catalysis and is degraded into pyruvate and acetaldehyde, which are precursor molecules for energy production in the cell.

Pangenome analysis of the bacterial genomes revealed 13,897 homologous gene families (Figure 5). The positive isolate contributed 3,555 genes to these families, with 222 unique genes. The observation that genes required for degrading PAHs share a common ancestry suggests that hydrocarbon-degrading genes are largely homologous, with only a few exceptions.

**Figure 5.**
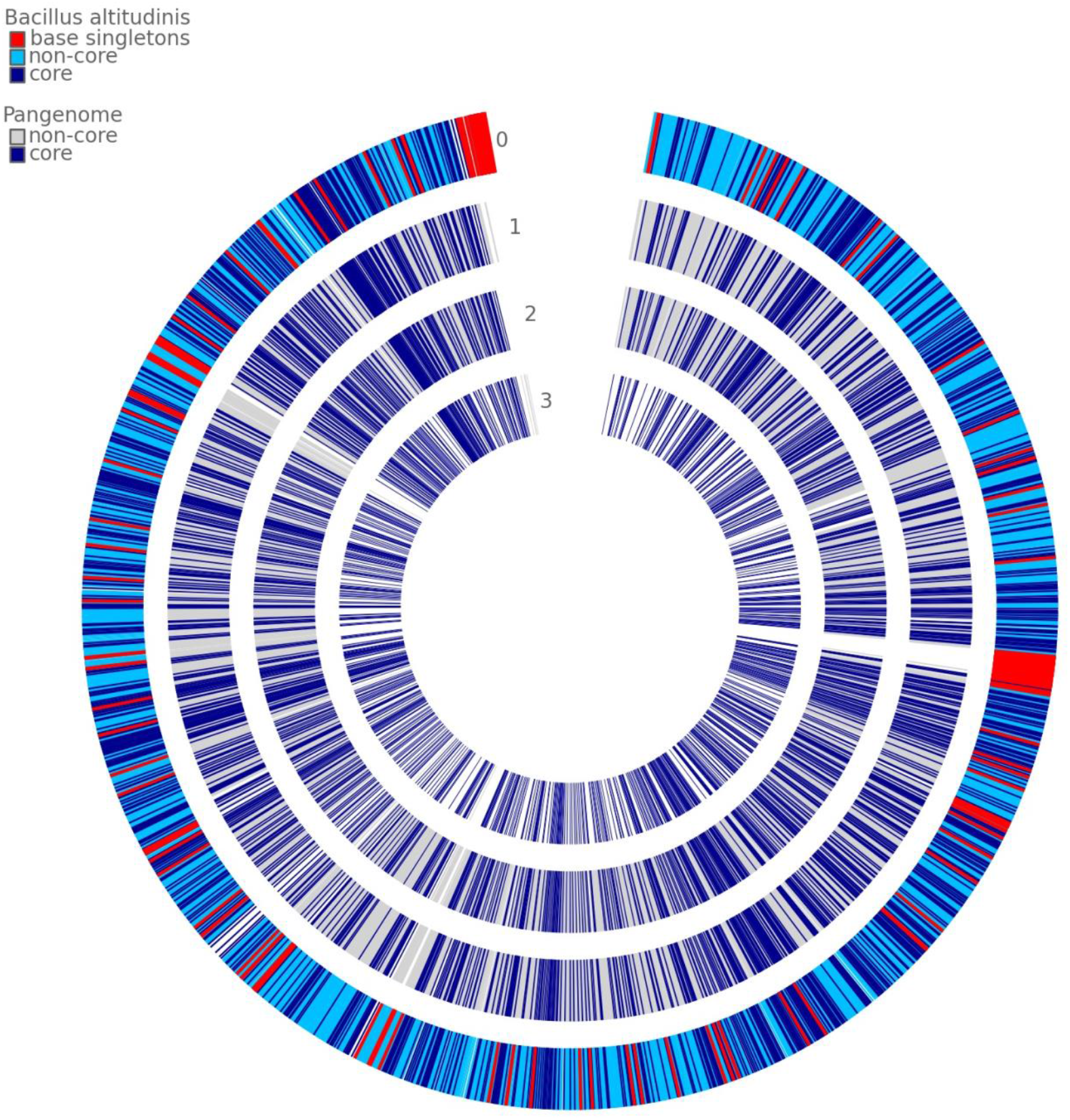
Pangenome circle plot of Bacillus altitudinis, Bacillus pumilus, and Mycolicibacterium vanbaalenii PYR-1. AR19 was assigned as 0 to represent the core genome. Singleton genes are highlighted in red.

From the genomic data of AR19, it can be hypothesized that this hydrocarbon-degrading isolate possesses multiple mechanisms that facilitate its survival in harsh environments. However, it remains unclear whether these mechanisms are universal to all bacteria or are specific to benzo[a]pyrene-degrading bacteria. Therefore, future research should aim to elucidate the unique characteristics of hydrocarbon-degrading bacteria by investigating all the cellular mechanisms responsible for protection against hydrocarbon stress.

## Data availability

The sequencing data have been deposited in the NCBI database The raw reads and assembly can be accessed using the accession numbers SRP540094 and QAWE00000000, respectively.

## Acknowledgment

The sequencing of the genome was funded by CSIR-INDEPTH (BSC-0111) network. Dr. Dasgupta would like to acknowledge the Director of CSIR-NEIST (India), Vice Chancellor of Dibrugarh University (India) and Dr. Ratul Saikia (CSIR-NEIST, India) (PI of BSC-0111) for providing facilities to conduct the research. Dr. Dasgupta would also like to acknowledge the chairperson of CBB, Dibrugarh University (India).

## Author contribution

ADG conceptualized the study, prepared the original draft of the manuscript, and contributed to review and editing. BN performed the flow cytometry experiments. RS performed the biosurfactant study.

## References

1. Zhang W, Mao G, Zhuang J, Yang H. The co-inoculation of Pseudomonas chlororaphis H1 and Bacillus altitudinis Y1 promoted soybean [Glycine max (L.) Merrill] growth and increased the relative abundance of beneficial microorganisms in rhizosphere and root. Front Microbiol [Internet]. 2023 Jan 9 [cited 2025 Sep 6];13. Available from: https://www.frontiersin.org/journals/microbiology/articles/10.3389/fmicb.2022.1079348/full

2. Kaushal P, Pati AM. Bacillus altitudinis Mediated Lead Bioremediation for Enhanced Growth of Rice Seedlings. Curr Microbiol. 2024 Oct 16;81(12):410.

3. Shivaji S, Chaturvedi P, Suresh K, Reddy GSN, Dutt CBS, Wainwright M, et al. Bacillus aerius sp. nov., Bacillus aerophilus sp. nov., Bacillus stratosphericus sp. nov. and Bacillus altitudinis sp. nov., isolated from cryogenic tubes used for collecting air samples from high altitudes. Int J Syst Evol Microbiol. 2006 Jul;56(Pt 7):1465–73.

4. Yue Z, Chen Y, Wang Y, Zheng L, Zhang Q, Liu Y, et al. Halotolerant Bacillus altitudinis WR10 improves salt tolerance in wheat via a multi-level mechanism. Front Plant Sci [Internet]. 2022 Jul 14 [cited 2025 Sep 6];13. Available from: https://www.frontiersin.org/journals/plant-science/articles/10.3389/fpls.2022.941388/full

5. Tai H, Guo Q, Zhao J, Liu Y, Yu H, Liu Y, et al. A thermostable xylanase hydrolyzes several polysaccharides from Bacillus altitudinis JYY-02 showing promise for industrial applications. Carbohydr Res. 2024 Apr 1;538:109080.

6. Ma XH, Shen S, Li W, Wang J. Bioherbicidal potential of Bacillus altitudinis D30202 on Avena fatua L.: a whole-genome sequencing analysis. J Appl Genet. 2023 Dec;64(4):809–17.

7. Venkatraman G, Giribabu N, Mohan PS, Muttiah B, Govindarajan VK, Alagiri M, et al. Environmental impact and human health effects of polycyclic aromatic hydrocarbons and remedial strategies: A detailed review. Chemosphere. 2024 Mar 1;351:141227.

8. Abdel-Shafy HI, Mansour MSM. A review on polycyclic aromatic hydrocarbons: Source, environmental impact, effect on human health and remediation. Egypt J Pet. 2016 Mar 1;25(1):107–23.

9. Bukowska B, Mokra K, Michałowicz J. Benzo[a]pyrene—Environmental Occurrence, Human Exposure, and Mechanisms of Toxicity. Int J Mol Sci. 2022 Jun 6;23(11):6348.

10. Steffen KT, Hatakka A, Hofrichter M. Degradation of Benzo[a]pyrene by the Litter-Decomposing Basidiomycete Stropharia coronilla: Role of Manganese Peroxidase. Appl Environ Microbiol. 2003 Jul;69(7):3957–64.

11. Serial Dilution Protocols [Internet]. [cited 2025 Sep 6]. Serial Dilution Protocols. Available from: https://asm.org:443/protocols/serial-dilution-protocols

12. Zhao B, Wang H, Mao X, Li R. A rapid screening method for bacteria degrading polycyclic aromatic hydrocarbons. Lett Appl Microbiol. 2009 Sep;49(3):408–10.

13. Nayarisseri A, Singh P, Singh SK. Screening, isolation and characterization of biosurfactant producing Bacillus subtilis strain ANSKLAB03. Bioinformation. 2018 Jun 30;14(6):304–14.

14. Dasgupta A, Saikia R, Kakoti BB, Handique PJ. Draft genome sequence of benzo[a]pyrene degrading Bacillus altitudinis strain AR19 isolated from Digboi oil refinery (India). Microbiol Resour Announc. 2025 Mar 13;14(4):e00957–24.

15. GTDB-Tk: a toolkit to classify genomes with the Genome Taxonomy Database | Bioinformatics | Oxford Academic [Internet]. [cited 2025 Sep 6]. Available from: https://academic.oup.com/bioinformatics/article/36/6/1925/5626182

16. Li L, Stoeckert CJ, Roos DS. OrthoMCL: identification of ortholog groups for eukaryotic genomes. Genome Res. 2003 Sep;13(9):2178–89.

17. Bhaganna P, Volkers RJM, Bell ANW, Kluge K, Timson DJ, McGrath JW, et al. Hydrophobic substances induce water stress in microbial cells. Microb Biotechnol. 2010 Nov;3(6):701–16.

18. Moe PC, Blount P, Kung C. Functional and structural conservation in the mechanosensitive channel MscL implicates elements crucial for mechanosensation. Mol Microbiol. 1998 May;28(3):583–92.

19. Whatmore AM, Chudek JA, Reed RH. The effects of osmotic upshock on the intracellular solute pools of Bacillus subtilis. J Gen Microbiol. 1990 Dec;136(12):2527–35.

20. Boch J, Kempf B, Bremer E. Osmoregulation in Bacillus subtilis: synthesis of the osmoprotectant glycine betaine from exogenously provided choline. J Bacteriol. 1994 Sep;176(17):5364–71.

21. Kempf B, Bremer E. OpuA, an osmotically regulated binding protein-dependent transport system for the osmoprotectant glycine betaine in Bacillus subtilis. J Biol Chem. 1995 Jul 14;270(28):16701–13.

22. Pathak KV, Keharia H. Application of extracellular lipopeptide biosurfactant produced by endophytic Bacillus subtilis K1 isolated from aerial roots of banyan (Ficus benghalensis) in microbially enhanced oil recovery (MEOR). 3 Biotech. 2014 Feb;4(1):41–8.

23. Omarova M, Swientoniewski L, Tsengam IM, Blake D, John V, McCormick A, et al. Biofilm Formation by Hydrocarbon-Degrading Marine Bacteria and Its Effects on Oil Dispersion. ACS Sustain Chem Eng. 2019 Sep 3;7(17):14490–9.

24. Xie Y, Yu F, Wang Q, Gu X, Chen W. Cloning of catechol 2, 3-dioxygenase gene and construction of a stable genetically engineered strain for degrading crude oil. Indian J Microbiol. 2014 Mar;54(1):59–64.

